# Syndecan-1 facilitates breast cancer metastasis to the brain

**DOI:** 10.1101/565648

**Authors:** Megan R. Sayyad, Madhavi Puchalapalli, Natasha G. Vergara, Sierra Mosticone Wangensteen, Melvin Moore, Liang Mu, Chevaunne Edwards, Aubree Anderson, Stefanie Kall, Megan Sullivan, Mikhail Dozmorov, Jaime Singh, Michael O. Idowu, Jennifer E. Koblinski

## Abstract

**Purpose:** Although survival rates for patients with localized breast cancer have increased, patients with metastatic breast cancer still have poor prognosis. Understanding key factors involved in promoting breast cancer metastasis is imperative for better treatments. In this study, we investigated the role of syndecan-1 (Sdc1) in breast cancer metastasis.

**Methods:** To assess the role of Sdc1 in breast cancer metastasis, we silenced Sdc1 expression in the triple-negative breast cancer human MDA-MB-231 cell line and overexpressed it in the mouse mammary carcinoma 4T1 cell line. Intracardiac injections were performed in an experimental mouse metastasis model using both cell lines. In vitro transwell blood-brain barrier (BBB) and brain section adhesion assays were utilized to specifically investigate how Sdc1 promotes brain metastasis. A cytokine array was performed to evaluate differences in the breast cancer cell secretome when Sdc1 is silenced.

**Results:** Silencing expression of Sdc1 in breast cancer cells significantly reduced metastasis to the brain. Conversely, overexpression of Sdc1 increased metastasis to the brain. We found that silencing of Sdc1 expression had no effect on attachment of breast cancer cells to brain endothelial cells or astrocytes, but migration across the BBB was reduced as well as adhesion to the perivascular regions of the brain. Loss of Sdc1 also led to changes in breast cancer cell-secreted cytokines/chemokines, which may influence the BBB.

**Conclusions:** Taken together, our study demonstrates a role for Sdc1 in promoting breast cancer metastasis to the brain. These findings suggest that Sdc1 supports breast cancer cell migration across the BBB through regulation of cytokines, which may modulate the BBB. Further elucidating this mechanism will allow for the development of therapeutic strategies to combat brain metastasis.

## Introduction

The five-year disease-free survival for breast cancer (BC) patients diagnosed with localized disease is 93-98%; however, for patients with distant-stage BC, e.g., brain metastasis, the survival is only 26-40% [1]. Women with brain metastases survive on average 2-16 months after diagnosis [2, 3]. Syndecan-1 (Sdc1) has been implicated in promoting breast cancer progression and is highly expressed in HER2+ and triple-negative breast cancers (TNBC) [4, 5]. Patients with TNBC or HER2+ breast cancer have a worse overall survival that is associated with increased metastatic burden [6].

Syndecans (Sdcs) are cell surface heparan sulfate proteoglycans (HSPGs) that act as co-receptors for growth factors, matrix proteins, cytokines and chemokines. Through these interactions, the Sdcs mediate several biological functions, such as cell-cell and cell-matrix adhesion, proliferation, differentiation, migration, and angiogenesis. Many have reported that the Sdcs play a dynamic role in cancer, whereby they can either promote or suppress cancer depending on the type of cancer and the type of Sdcs expressed[7–13]. Sdc1, typically expressed in epithelial cells, is widely detected in epithelial neoplasms derived from numerous origins [11, 14]. It is specifically associated with poor prognosis in BC patients, and it has been linked to rapid BC growth rate [8, 15]. Up-regulation of mRNA and protein levels of Sdc1 have been detected in BC cell lines and patient biospecimens [4, 16]. Further, high levels of Sdc1 correlate with lower overall survival (OS), relapse-free survival (RFS), distant metastasis-free survival (DMFS) and worse metastatic relapse-free survival (MRFS) in BC patients [4, 7, 8, 17, 18]. Strong stromal expression of Sdc1 is a negative prognostic factor in BC and correlates with poor response to chemotherapy [11]. Taken together, these observations indicate that Sdc1 is associated with higher tumor grade, malignant forms of BC, and poor clinical outcome in patients [10].

Several in vitro and in vivo studies are congruent with the clinical expression of Sdc1 in BC. For instance, loss of Sdc1 suppresses Wnt-1-induced mammary carcinoma [19]. Moreover, Sdc1 is observed in stromal fibroblast and endothelial cells of invasive breast carcinomas where it is involved in tumor growth and angiogenesis [20]. Finally, Sdc1 directly participates in tumor cell spreading and adhesion, and through interaction with integrins, it promotes invasion [21, 22]. Notwithstanding this progress in understanding the diverse pro-malignant functions of Sdc1, little is known about its role in BC metastasis.

In the present study, we found that Sdc1 is important in mediating metastasis to the brain. Our findings indicate that Sdc1 acts by facilitating transmigration of BC cells through the BBB, but has no effect on attachment of BC cells to brain endothelial cells or astrocytes. Thus, we explored how Sdc1 promotes this migration, and found that it affects secretion of a variety of cytokines/chemokines, many of which have been implicated in vascular permeability and cell migration. As such, our findings provide insight into how BC cells can invade and grow in the unique brain environment and the role that Sdc1 plays in this process. We believe this study will help with future development of therapeutic strategies to combat BC metastasis to the brain.

## Materials and methods

### Cell culture and reagents

The human TNBC cell line, MDA-MB-231 (MDA-231; gift from Dr. D. Welch, University of Kansas Cancer Center), was grown in DMEM/F-12 (ThermoFisher) supplemented with 5% FBS, L-glutamine (2mM), sodium pyruvate (1mM), and non-essential amino acids (0.02mM, Gemini Bio-Products). The mouse mammary carcinoma cell line 4T1 (gift from Dr. S Spiegel, Virginia Commonwealth University [VCU]) was grown in RPMI 1640 (ThermoFisher) supplemented with 10% FBS (Gemini Bio-Products). BT549 cells were cultured in RPMI 1640 supplemented with 10% FBS and insulin (1ug/ml). Immortalized human umbilical vein endothelial cells (iHUVEC, gift from Dr. W. Muller, Northwestern University [NU]) were cultured in human endothelial serum-free medium (Gibco) supplemented with 10% FBS, endothelial cell growth supplement (0.01mg/ml, Corning), and L-glutamine (2mM). Human cerebral microvascular endothelial cells (hCMEC/D3 [hCMEC]; gift from Dr. B. Weksler, Cornell Medical College) were grown in endothelial basal medium-2 (Lonza) supplemented with 5% FBS, basic fibroblast growth factor (1ng/ml), hydrocortisone (1.4mM), ascorbic acid (5mg/ml) and chemically defined lipid concentrate (1/100 dilution, ThermoFisher). All cell media contained penicillin (100U/ml) and streptomycin (100µg/ml) (ThermoFisher). Primary cortical human astrocytes (HA) were grown on poly-l-lysine (0.02mg/ml, Bio-Techne) coated dishes with HA media kit (ScienCell). All cells were maintained at 37°C in a 5% CO_2_/95% humidified air atmosphere and were routinely checked for mycoplasma. Cell lines were validated by Short Tandem Repeat analysis (DNA Diagnostic Center).

### Invasion and migration assays using the xCELLigence RTCA DP System

Invasion and migration assays were performed using 8µm-pore CIM (Cell Invasion/Migration)-Plate in the xCELLigence Real-Time Cell Analysis System (ACEA Biosciences) according to manufacturer’s recommendations. For invasion assays, the upper chambers of the CIM-plate were pre-coated for 4 hours with phenol red-free basement membrane extract (BME/Matrigel) (0.6mg/mL) at 37°C. MDA-231 NS1 and Sdc1 KD cell (100,000/well for migration; 20,000/well for invasion; seeded in 5% FBS-containing media) migration/invasion towards 20% FBS-containing media was measured for 24 hours. The data was normalized in the xCELLigence software and results are reported as Mean Delta Normalized Cell Index (CI). High CI values are indicative of increased cell invasion and migration. These experiments were completed three times in quadruplicate. Statistical analysis, two-way ANOVA with Sidak’s multiple comparisons was performed.

### Animal experiments

All animal experiments were conducted in accordance with a protocol approved by NU and VCU Institutional Animal Care and Use Committee. For all injections, cell lines were collected using Versene (EDTA, ThermoFisher). The cells were mixed with phenol red-free BME (14.5mg/ml) for intraductal injections. The MDA-231 cell/BME mixture (100 µl) was injected into the lactiferous duct of the fourth mammary glands on both sides of either 6-week-old female athymic nude (0.5×10^6^ cells/gland, Foxn1^nu/nu^, Envigo) or NSG mice (0.25×10^6^ cells/gland, NOD.Cg-*Prkdc^scid^ Il2rg^tm1Wjl^*/SzJ, Jackson Labs) as described [23, 24]. Tumors were measured biweekly beginning 21 days after injection, and tumor area was calculated by width x length (mm^2^). Entire primary mammary tumors were removed and weighed. Additionally, MDA-231 (2×10^5^) or 4T1 (1×10^5^) cells were injected into the left ventricle of the heart of 4-week-old nude, NSG, or Balb/cJ mice, respectively, as described [25]. Animals were sacrificed 21 days after injections. For both intraductal and intracardiac injections, all visceral organs, bones, brain, and lymph nodes were harvested for ex vivo examination of metastasis. To examine growth in the brain, the MDA-231 cells (1×10^4^ cell/5 µl) were directly injected into the brain (0 Bregma, right 2.5 mm lateral, 3 mm deep) as described [26]. For all studies, visible metastases were imaged and counted using a Zeiss Stemi SV11 Apo fluorescent dissecting microscope. Metamorph analysis software was used to quantify brain metastases in cranial and caudal images. Statistics were determined by Student’s unpaired t-test.

### In vitro transwell BBB assay

An in vitro BBB model was constructed as previously described [27]. Briefly, iHUVEC were co-cultured with HA on opposite sides of an 8µm-pore, PET FluoroBlok membrane transwell insert (BD Biosciences) for 3 days. Astrocytes (1×10^5^ cells) were plated on the bottom side (day 1) and endothelial cells (5×10^4^ cells) were plated on the upper side (day 2) of the membrane. Before seeding the cells, the membrane was coated with poly-l-lysine (HA, 0.02mg/ml, Bio-Techne) and gelatin (iHUVEC, 0.2%, Sigma). hCMEC were used in a similar format. After 2 days of iHUVEC/HA growth together, BC cells (1×10^5^/well) were seeded on iHUVECs and migration towards BBB medium was assessed after 24 hours [27]. Following fixation (4% PFA for 20 min), cells were stained with DAPI (167ng/ml). The number of migrated tumor cells was analyzed from 6 images/well (Zeiss AxioVert 100 fluorescent microscope) using CellProfiler software. Statistics were determined by one-way ANOVA with Bonferroni multiple comparison post-test.

### In vitro brain section adhesion assay

The in vitro tissue section assay was performed as described [28]. Tumor cells were seeded onto fresh-frozen mouse brain (coronal), lung, or liver 20 µm sections. After 90 minutes, cells were fixed in 4% PFA, DAPI-stained, cover-slipped with Prolong Gold mounting medium (ThermoFisher) and imaged using fluorescence microscopy. The number of tumor cells attached to each section was analyzed using ImageJ software. Statistics were determined by Student’s unpaired t-test or one-way ANOVA with Bonferroni multiple comparison post-test.

### Clinical samples and immunohistochemistry

Primary human breast tumors and BC brain metastases were obtained from the NU and VCU Pathology departments. Tissue microarrays (TMA) were made from 7 matched primary tumor and brain metastases from each institution and a total of 63 unmatched brain metastases (77 total). The tissue was obtained in compliance with protocols approved by the NU and VCU’s Institutional Review Boards. Immunohistochemistry for Sdc1 (1:50 dilution of B-A38, Bio-Rad) was performed by the VCU Cancer Mouse Models Core Laboratory with the Leica Bond RX auto-stainer using heat-induced epitope retrieval buffer 1 (Leica, Sodium Citrate buffer, pH 6.0). Stained slides were then imaged on the Vectra Polaris (Akoya Biosciences). Immunoreactivity staining was evaluated and scored by clinical pathologists (MOI, JS, MS) in a blinded fashion.

TCGA level 3 gene expression data summarized as RSEM values was obtained using the TCGA2STAT R package v 1.2, along with the corresponding clinical annotations. The data was log2-transformed and analyzed using Kaplan-Meier curves and Cox proportional hazard model. Hazard ratio with 95% confidence intervals and logrank P value were reported. Each gene of interest was analyzed for its effect on survival by separating patients into high/low expression subgroups. A modified approach from was used to estimate the best gene expression cutoff that separates high/low expression subgroups with differential survival [29]. The survival analysis was performed in R/Bioconductor statistical environment v.3.5.1 [30].

## Results

### Silencing of Sdc1 expression has no effect on cell proliferation and invasion in vitro

To investigate the role of Sdc1 in BC, we silenced its expression in MDA-231 cells (Sdc1 KD; 75% decrease both at the message and protein levels as compared to the NS1 (control expressing non-silencing shRNAmir). Of note, the other Sdcs were not significantly changed in Sdc1 KD cells [31]. For all experiments, Versene was used to collect cells to avoid cleavage of HSPGs from the cell surface, which occurs with trypsin [32]. Silencing of Sdc1 in MDA-231 cells had no effect on cell proliferation compared to NS1 cells irrespective of whether the cells were grown in serum or serum-free media (Fig. 1a; Supplementary Fig. 1a). Others have shown that loss of Sdc1 resulted in increased cell migration in keratinocytes and MDA-231 cells [33–35]. Indeed, in MDA-231 Sdc1 KD cells, we observed increased migration compared to NS1 cells using the xCELLigence System (Fig. 1b). However, there was no change in invasion through BME/Matrigel in the xCELLigence System (Fig. 1b). Similar results were observed using traditional transwell systems (data not shown). In conclusion, these experiments confirmed that loss of Sdc1 has no effect on MDA-231 cell proliferation and invasion, and only a slight increase in cell migration.

**Figure 1.**
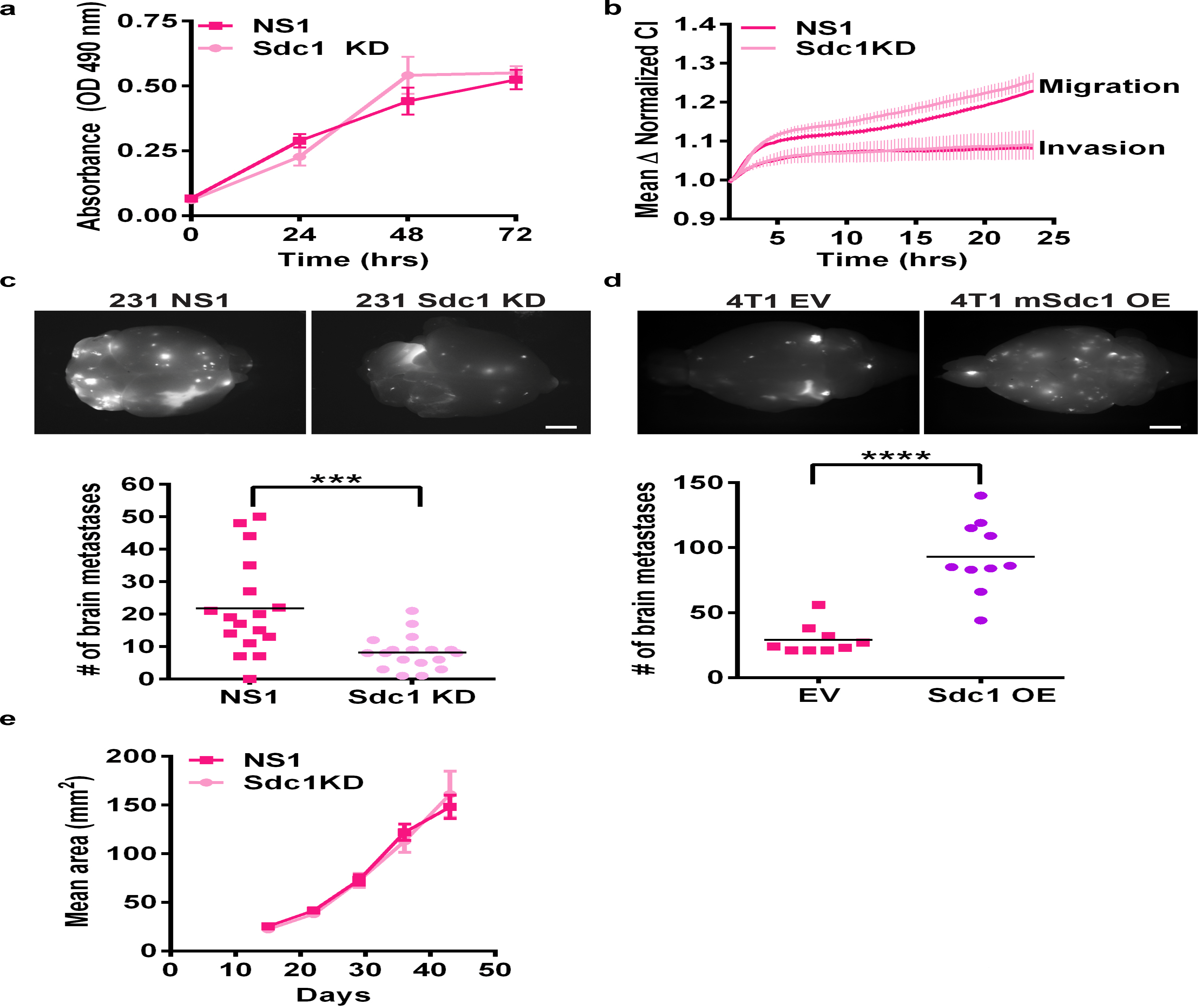
Sdc1 expression facilitates breast cancer metastasis to the brain. **a** Silencing of Sdc1 expression does not affect proliferation of MDA-231 cells grown in serum-free media on BME/Matrigel. **b** Silencing of Sdc1 expression in MDA-231 cells had no significant effect on cell migration or invasion in vitro compared to MDA-231 NS1 cells. n=3. CI, Cell Index. **c** Top, representative fluorescent images of the cranial side of the brain after intracardiac injection of MDA-231 NS1 and Sdc1 KD cells in nude mice. Bottom, Mice injected with MDA-231 Sdc1 KD cells had a significant reduction in the number of metastases on the surface of the brain. NS1, n=17, Sdc1 KD, n=18. ***p<0.001, Similar results were seen in NSG mice (data not shown). **d** Top, representative fluorescent images of the cranial side of the brain after intracardiac injection of control 4T1 EV and mSdc1 OE mammary cancer cells in Balb/cJ mice. Bottom, mice injected with 4T1 mSdc1 OE cells had a significant increase in metastases on the surface of the brain. EV, n=10, Sdc1 OE, n=18. ****p<0.0001. **e** The growth rate of primary tumors arising from MDA-231 cells with reduced expression of Sdc1 was not significantly different from the control (NS1) cell lines, NSG mice. NS1, n=19, Sdc1 KD, n=18. Similar results were seen in nude mice (data not shown). For **a, b, e** Data point, mean ± SEM. **c & d** Top, Bar, 2mm.

### Sdc1 mediates BC metastasis in vivo

To determine if Sdc1 has effects on BC primary tumor growth and metastasis in vivo, we injected both MDA-231 NS1 and Sdc1 KD cells into the fourth mammary fat pad of mice to allow for the formation of spontaneous tumors. After approximately 60 days, the mice were sacrificed and the lymph node (LN), lungs, liver, brain, bone, ovary and heart were collected to determine the percent of mice with metastases to these organs (Table 1). The primary tumor growth was not significantly different (Fig 1e). Of all the organs assessed, differences in metastases following MDA-231 NS1 and Sdc1 KD cell injection were observed in only the liver (NS1: 37% v Sdc1 KD: 17%) and the brain (NS1: 53% v Sdc1 KD: 33%); however, these differences were not significant. Additionally, no differences were observed in other organs (Table 1). We hypothesized that the brain and bone metastases did not have sufficient time to grow out due to extensive lung metastases. Thus, we performed intracardiac injections (an experimental mouse model of metastasis) of MDA-231 NS1 and Sdc1 KD cells. Here, a significant decrease in the number of brain metastases was found in mice injected with MDA-231 Sdc1 KD cells as compared to mice injected with MDA-231 NS1 cells (Fig.1c). No significant difference in metastasis to other organs was detected (bone, lung, liver; Supplementary Fig. 1b-d). To determine whether this effect could be seen in an immune competent animal, we used an allograft mouse model. We overexpressed Sdc1 in the 4T1 mouse mammary carcinoma cell line (Supplementary Fig. 2a) and intracardially injected these cells or EV (empty vector) control cells into the syngeneic BALB/cJ mice. We found that the gain of Sdc1 expression resulted in significantly increased metastasis to the brain, bones, and liver (Fig. 1d; Supplementary Fig. 1e-g). These findings indicate that Sdc1 in tumor cells has no effect on primary tumor growth but plays a significant role in metastasis to specific organs, especially to the brain. We chose to focus on brain metastasis since a significant difference was observed in both the MDA-231 and 4T1 experimental metastasis models.

**Table 1.**
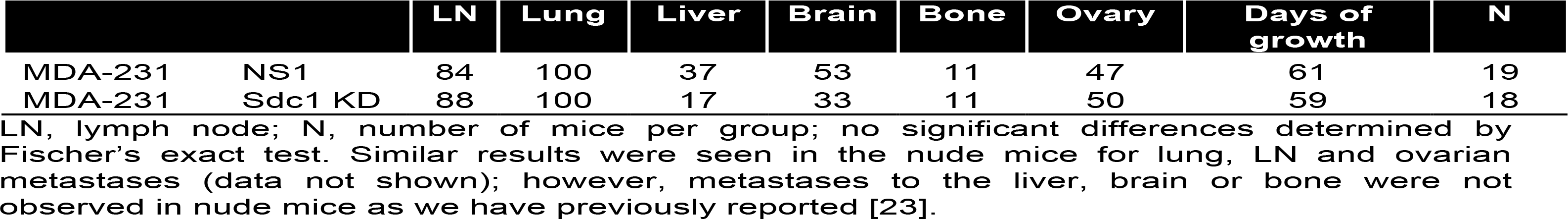
Percent of mice (NSG) with metastases from the mammary gland to specific organs.

|  |  | LN | Lung | Liver | Brain | Bone | Ovary | Days of growth | N |
| --- | --- | --- | --- | --- | --- | --- | --- | --- | --- |
| MDA-231 | NS1 | 84 | 100 | 37 | 53 | 11 | 47 | 61 | 19 |
| MDA-231 | Sdc1 KD | 88 | 100 | 17 | 33 | 11 | 50 | 59 | 18 |

LN, lymph node; N, number of mice per group; no significant differences determined by Fischer's exact test. Similar results were seen in the nude mice for lung, LN and ovarian metastases (data not shown); however, metastases to the liver, brain or bone were not observed in nude mice as we have previously reported [23].

**Figure 2.**
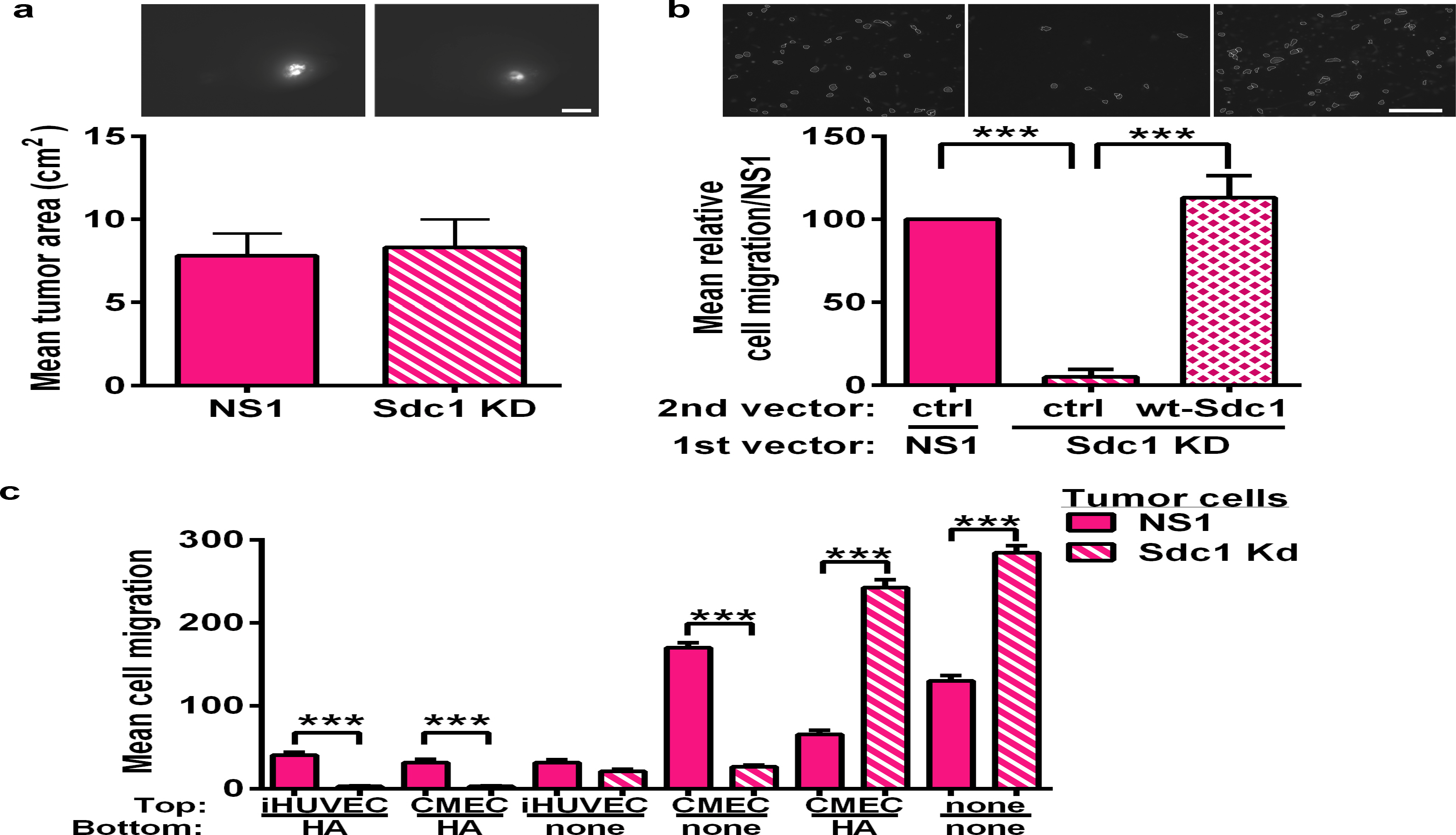
Sdc1 facilitates MDA-231 cell migration across the BBB. **a** Silencing of Sdc1 expression has no effect on BC growth within the brain when the cells were directly injected into the brain (intracranial injection). Top, fluorescent images of the cranial of the brain. Bottom, The mean tumor area ± SEM was not different in mice injected with MDA-231 Sdc1 KD cells compared to those injected with MDA-231 NS1 cells, n=4 for each group. **b** Representative images (top) and quantification (bottom) showing that silencing Sdc1 in MDA-231 cells results in decreased migration across an in vitro transwell BBB model system compared to NS1 control cells and MDA-231 Sdc1 KD cells expressing wt-Sdc1. The cells were transfected with either NS1 or Sdc1 shRNAmir (1^st^ vector) and then were infected with either empty (ctrl) vector or wt-Sdc1 (2^nd^ vector). Quantification of cell migration, n=4, ***p<0.001. **c** Silencing of Sdc1 expression decreases migration of the MDA-231 cells only when the cells are migrating through either hCMEC (CMEC) or a BBB made of iHUVEC and HA or CMEC and HA. In contrast, silencing Sdc1 expression increases migration of MDA-231 cells across HA alone or no cells (none) compared to control NS1 cells. n=3, ***p<0.001. For all experiments, Bars, mean ± SEM. **a & b** Top, Bar, 2mm.

### Sdc1 facilitates BC cell transmigration across the BBB and attachment to perivascular regions in the brain

To examine whether Sdc1 affects tumor growth after seeding to the brain, mice were intracranially injected with MDA-231 NS1 and Sdc1 KD cells. No difference was found in the size of brain tumors detected (Fig. 2a). This suggests that Sdc1 may affect the ability of BC cells to extravasate and migrate into the brain environment, rather than to facilitate tumor outgrowth in the brain.

We next assessed the role of Sdc1 in BC cell transmigration across the BBB using a previously established in vitro BBB transwell model [27]. As confirmed in this model previously, we saw up-regulation of the glucose transporter 1 and γ-glutamyl transpeptidase 1 on the endothelial iHUVEC cells when co-cultured with HA, suggesting that iHUVECs express BBB specific proteins (data not shown) [27]. Furthermore, tight junction formation was observed by prominent ZO-1 staining (data not shown) [27]. We also used hCMEC brain microvascular endothelial cells in this same model (Fig. 2c) [27, 36]. When Sdc1 was silenced, there was a significant reduction in transmigration of the MDA-231 cells only across the in vitro BBB composed of iHUVEC/HA, hCMEC/HA, or hCMEC alone compared to NS1 controls (Fig. 2b, c). Importantly, wild-type Sdc1 (wt-Sdc1) can rescue the reduced BBB transmigration seen with Sdc1 KD cells (Fig. 2b). To confirm that the reduction in BBB transmigration was not due to inability of the Sdc1 KD cells to adhere to the in vitro BBB, we seeded MDA-231 NS1 and Sdc1 KD cells onto a monolayer of either iHUVECs, hCMECs, or HAs, and observed no difference in the attachment to these cells (Supplementary Fig. 3). Additionally, MDA-231 Sdc1 KD cells exhibited significantly more migration than NS1 controls through the insert alone as well as across HAs alone (Fig. 2c). This is similar to our xCELLigence data (Fig. 1b) and previous reports [35]. Taken together, these findings support the notion that Sdc1 facilitates BC cell transmigration across the BBB.

**Figure 3.**
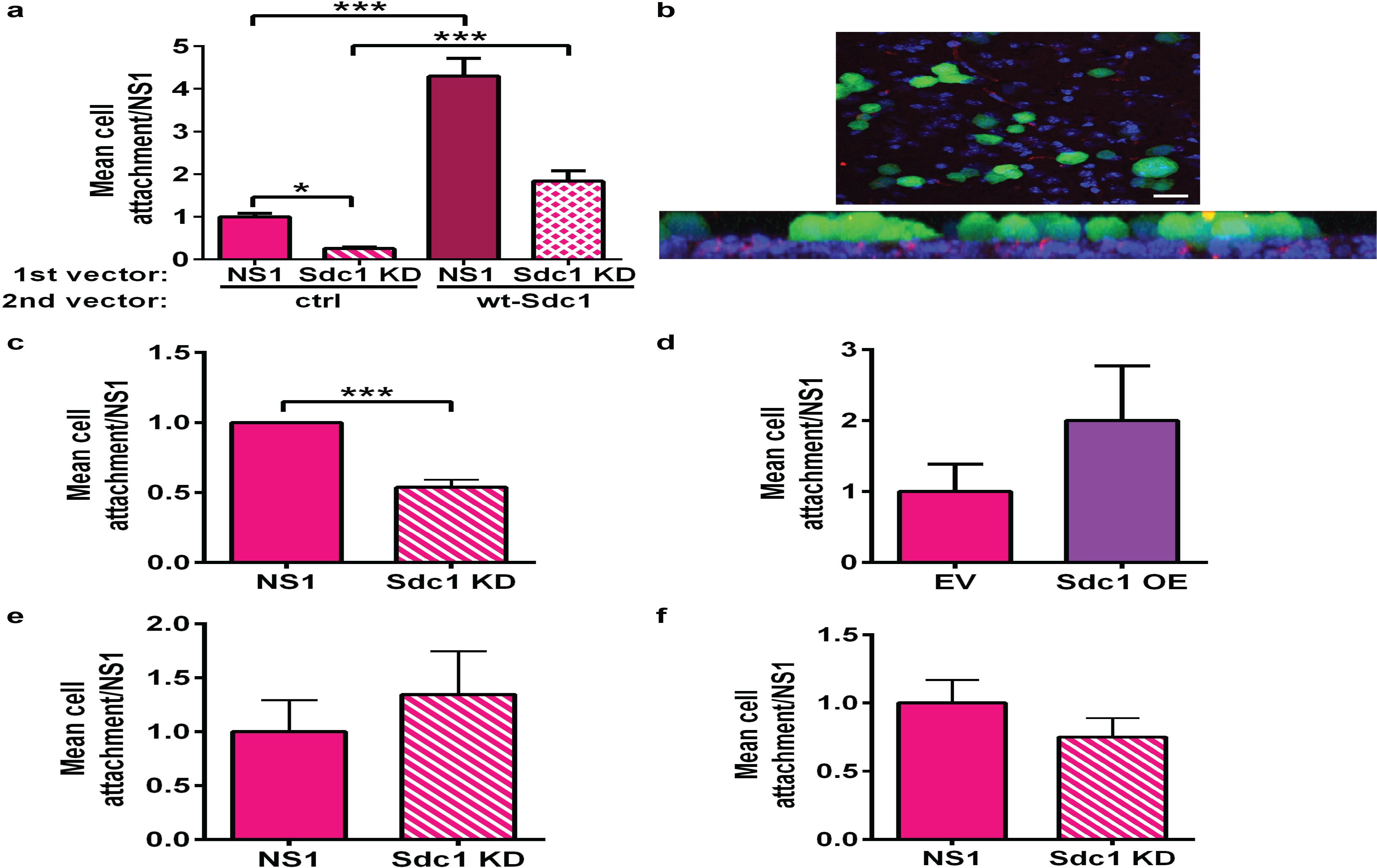
Sdc1 facilitates breast cancer cell attachment to the perivascular regions of the frozen brain section. **a** Overexpression of wt-Sdc1 in MDA-231 NS1 control cells increased the ability of the cells to bind to a frozen mouse brain section, and expression of wt-Sdc1 in Sdc1 KD cells rescued the ability for those cells to attach to the brain section. The cells were transfected with either NS1 or Sdc1 shRNAmir (1^st^ vector) and then were infected with either empty (ctrl) vector or wt-Sdc1 (2^nd^ vector). n=3, ***p<0.001. **b** Representative image of MDA-231 NS1 GFP positive cells (green) attaching to perivascular regions (PECAM-1/CD31 positive staining for endothelial cells, red) in the brain (nuclei of brain cells DAPI stained, blue). Top panel, x-axis, Bar, 20µm. Bottom panel, z-axis of confocal image. Silencing of Sdc1 (Sdc1 KD) in BT549 BC cells (**c**) decreases attachment, and overexpression of mouse Sdc1 in 4T1 mammary cancer cells (**d**) increases attachment of these cells to perivascular regions in frozen mouse brain section. Loss of Sdc1 has no effect on tumor cell attachment to frozen mouse liver (**e**) or lung sections (**f**). For all, n=3, Bars, mean ± SEM. ***p<0.001.

Carbonell et al. demonstrate that upon entry into the brain, cancer cells must first co-opt the vasculature to obtain nutrients and oxygen from blood circulation for successful colonization [27, 28]. To further understand the ability of Sdc1 to support metastatic formation in the brain, we examined attachment to perivascular regions using an in vitro brain section adhesion assay [28]. MDA-231 NS1 cells exhibited significantly greater attachment compared to Sdc1 KD cells (Fig. 3a). This effect was rescued by expression of wt-Sdc1, and enhanced attachment was observed in NS1 cells with enforced expression of wt-Sdc1 (Fig. 3a). Additionally, immunofluorescence using PECAM-1 (CD31) to stain the brain vasculature demonstrated that cancer cells have a preference for attachment to the perivascular regions of the brain (Fig. 3b). The same brain adhesion assay was also performed with BT-549 BC cells with silenced Sdc1 expression (Supplementary Fig. 2b) and 4T1 cells to confirm our results in other cell lines. We found that BT549 NS1 cells had significantly greater attachment compared to Sdc1 KD cells (Fig. 3c). Conversely, 4T1 Sdc1 OE cells displayed greater attachment to the brain section compared to EV control cells (Fig. 3d). To confirm that this difference in cell adhesion is specific to the brain, we performed a similar experiment using liver and lung sections, and found no significant differences in attachment between MDA-231 NS1 and Sdc1 KD cells (Fig. 3e, f). Overall, loss of Sdc1 resulted in decreased attachment in an in vitro brain section adhesion assay, suggesting that Sdc1 may also facilitate brain metastatic formation through BC cell adhesion in the brain environment.

### Sdc1 regulates cytokines and chemokines secreted by BC cells

It is well established that secreted factors from cells promote BBB disruption to create a leakier barrier for cell entry into the brain [37–40]. Thus, we investigated whether loss of Sdc1 would drive changes to the BC secretome, which could conceivably result in increased BBB permeability and BC cell adhesion to the perivascular regions in the brain. We specifically focused on cytokines and chemokines since they have been implicated in vascular permeability, inflammation, and cell migration, including transmigration across the BBB [39, 41–47]. Additionally, Sdc1 is known to interact with cytokines/chemokines, *e.g.*, IL-6 and IL-8 [22, 48, 49]. Thus, to determine if Sdc1 affects BC cell cytokine/chemokine production and/or release, we performed a cytokine/chemokine array followed by a multiplex approach to confirm our results using 24-hr conditioned medium (CM) collected from MDA-231 NS1 and Sdc1 KD cells. We identified differences in secreted cytokine/chemokine levels with a significant difference in GRO-α, ICAM1, IL-6, IL-8, GM-CSF and CCL5 (Fig. 4; Supplementary Fig. 4). Taken together, these findings indicate that Sdc1 plays a vital role in regulating cytokine and chemokine production and/or secretion from BC cells.

**Figure 4.**
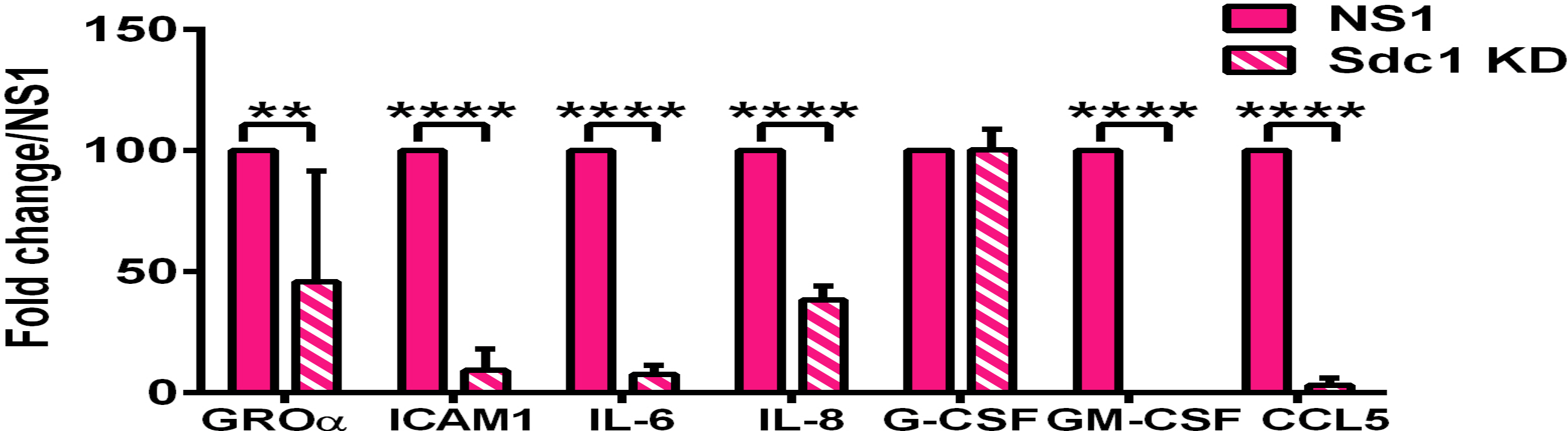
Silencing Sdc1 decreases cytokine/chemokine levels in MDA-231 cells. Quantification of human cytokines/chemokines from MDA-231 NS1 and Sdc1 KD cell 24-hr CM using a multiplex assay. **p<0.01, ****p<0.0001.

### BC patient brain metastases samples stained positive for Sdc1

To assess the clinical relevance of Sdc1 in BC and specifically in brain metastasis, we performed meta-analysis using The Cancer Genome Atlas (TCGA) and Kaplan-Meier Plotter [50]. TCGA analysis revealed a greater than 4-fold increase in Sdc1 expression in BC versus normal breast tissue (Fig. 5a). Using Kaplan-Meier analysis of the TCGA data, we found that regardless of BC subtype, patients with high Sdc1 expression had worse disease-free overall survival than those with low Sdc1 expression (Fig. 5b). Focusing on the TNBC subtype, which is associated with an increased risk for brain metastasis, we found that patients with high Sdc1 have significantly worse disease-free overall survival compared to those with low Sdc1 expression (Fig. 5c). We also examined brain metastases samples from BC patients for Sdc1 protein expression using a TMA. Tissue samples were scored from 0 to 8 on the Allred scale. Sixty-two out of 77 (81%) tissue samples presented positive staining for Sdc1 with a score of 3 or above (Fig. 5d-g). The majority of these brain metastases arose from patients with either TNBC or HER2+ tumors (Fig. 5h). These findings demonstrate the clinical relevance of Sdc1 in brain metastasis and indicate that high Sdc1 expression is associated with worse prognosis and brain metastatic formation.

**Figure 5.**
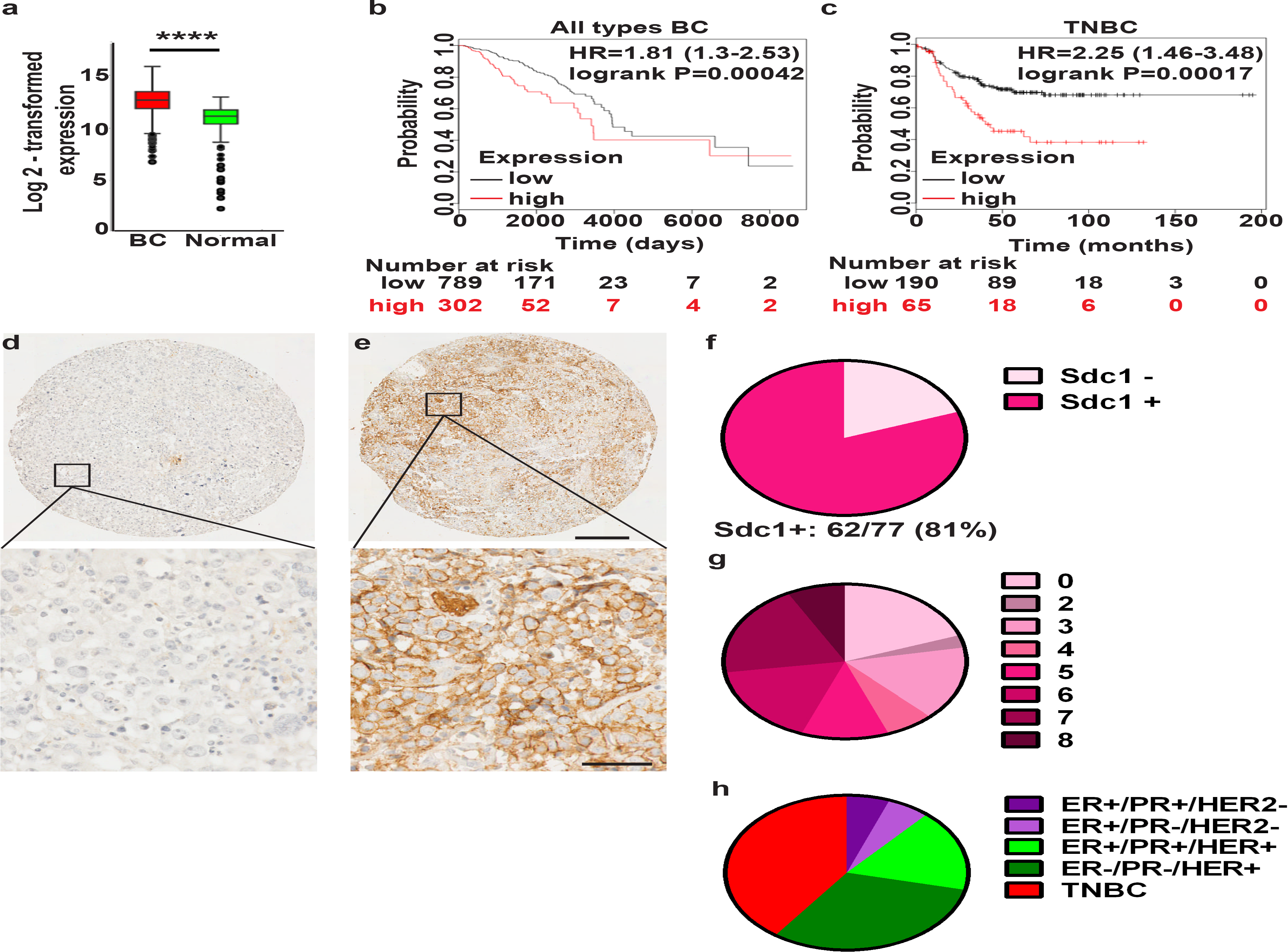
Sdc1 is expressed in human breast cancer patient brain metastases. **a** TCGA data showing expression of Sdc1 is significantly different in cancerous versus normal breast tissue. ****p<2E-16. **b** Disease-free overall survival of BC patients of all subtypes is significantly different between patients with high and low Sdc1 expression. n=1091, p=0.00042. **c** Disease-free overall survival of TNBC patients is significantly different between patients with high and low Sdc1 expression. n=255, p=0.00017. **d** Negative and **e** positive staining for Sdc1 in BC brain metastases TMA. Top, Bar, 200µm, Bottom, Bar, 50 µm. **f** Percentage of BC metastatic brain samples that stained positive or negative for Sdc1. **g** TMA Allred scores (0-2, negative and 3-8, positive) for 77 patients. No patients received a score of 1. **h** Proportion of primary tumor subtype (for which this information was available) for each brain metastases present in the TMA.

## Discussion

In this study, we set out to determine if Sdc1 plays a role in mediating BC metastasis. We found that alterations in Sdc1 expression had no effect on BC proliferation or invasion in vitro (Fig 1a, b). Additionally, changes in Sdc1 expression did not affect primary tumor growth (Fig. 1e). Although Sdc1 KD increases migration in vitro (Fig. 1b, 2c) [35], this phenomenon did not correlate with changes in overall metastasis in vivo. Interestingly, a more biologically complex in vitro migration assay to analyze the BBB correlated well with our in vivo data (Fig. 1c, 2b, c), in which we found that alterations in Sdc1 expression affect metastasis to the brain (Fig. 1c, d). BC patients with high Sdc1 expression have worse overall prognosis, and a high prevalence of BC patients’ brain metastases express Sdc1 (Fig. 5b-g) [4, 7, 10]. Intriguingly, others have demonstrated that high Sdc1 expression on stromal fibroblasts and endothelial cells supports primary BC growth and angiogenesis and allows for an invasive-permissive microenvironment, but this was model dependent [51–53]. Furthermore, Sdc1 expression on stromal fibroblasts is required for efficient BC metastasis to the lung using the 4T1 mouse mammary carcinoma model [54]. Taken together, these reports along with our findings presented here demonstrate the dynamic roles of Sdc1 in tumor cells, fibroblasts, and endothelium in promoting BC growth and metastasis.

Although a high number of BC patient’s brain metastases expressed Sdc1, the silencing of Sdc1 expression in BC did not affect growth of these cells when they were directly injected into the brain (Fig. 2a). These results suggest that the decrease in brain metastases seen when Sdc1 is silenced is due to Sdc1 facilitating arrival of BC cells to the brain and not growth in the brain. To seed the brain, metastatic cancer cells must cross a major obstacle, the BBB, a highly impenetrable physical and metabolic barrier composed of endothelial cells, astrocytes, and pericytes, and then bind to the perivascular regions in the brain [55, 56]. Using an in vitro transwell BBB model system, we found that Sdc1 facilitated BC cell transmigration across the BBB (Fig. 2b, c). Additionally, we found that Sdc1 plays an integral role in facilitating BC cell adhesion to the perivascular regions in the brain (Fig. 3a-d), but not to these regions in the liver and lungs (Fig. 3e, f). Interestingly, Sdc1 expression had no effect on BC cell binding to endothelial cells or astrocytes in vitro (Supplementary Fig.3). This suggests that Sdc1 affects motility of BC cells through the BBB and/or BBB permeability. Furthermore, it suggests that attachment of BC cells to perivascular regions in the brain occurs through binding of Sdc1 to the brain microenvironment. Alterations in Sdc1 expression had no effect on BC cell growth in the brain (Fig. 2a), indicating that Sdc1 specifically supports the efficient transit of cancer cells through the BBB.

Cancer cell migration across the BBB often involves disruption of it, particularly at the level of endothelial cell tight junctions as was shown for successful TNBC cell entry into the brain [37, 38, 40, 56, 57]. This disruption often occurs through secreted factors, e.g., cytokines/chemokines. Several groups have demonstrated that Sdc1 can regulate cytokines/chemokines, *e.g.*, IL-6, IL-8, IL-34 [22, 48, 58]. Thus, we explored the impact of Sdc1 on the BC cell secretome, focusing on cytokines/chemokines. Our cytokine study revealed that Sdc1 regulates cytokine/chemokine production and/or secretion from BC cells, as gauged by the significant decrease in these factors present in the CM collected from MDA-231 Sdc1 KD cells versus NS1 cells (Fig. 4, Supplementary Fig. 4). Many of the cytokines/chemokines we identified have previously been reported to be involved in regulating the blood-tumor-barrier, *e.g.,* GRO-α, ICAM-1, IL-6, IL-8, GM-CSF, and CCL5 [59]. Others have found that these proteins also have an impact on BBB integrity and/or facilitate BBB transmigration, *e.g.,* ICAM-1, GM-CSF, CCL5 [44–47]. Additionally, IL-8 was shown to bind to Sdc1, thereby facilitating leukocyte transendothelial migration [48]. Overall, these reports suggest that Sdc1 regulation of cytokines/chemokines is important in supporting cancer cell entry into the brain. It would be interesting to identify in the future the underlying mechanisms leading to brain metastasis in cells with high Sdc1 expression, and the key cytokine(s)/chemokine(s) involved in disrupting BBB homeostasis during breast cancer metastasis to the brain in vivo.

Here, we provide clinical evidence that high Sdc1 expression is associated with brain metastases and correlates with lower disease-free overall survival, especially in patients with TNBC (Fig. 5). Unraveling the mechanism for Sdc1-supported brain metastasis could guide therapy development. Zoledronate (FDA-approved drug for multiple myeloma and bone metastases) acts by targeting various extracellular matrix components, including Sdc1, to inhibit cancer progression [11, 60]. Radioimmunotherapy to target Sdc1 showed promising results in mice with TNBCs expressing Sdc1 [5]. Indeed, targeting tumors with high Sdc1 using a Sdc1-antibody-DM4 cytotoxic drug conjugate, Indatuximab ravtansine, exerts anti-tumor effects similar to trastuzumab emtansine. These results demonstrate that Sdc1 is a suitable target for TNBC therapy, and these treatment options could be explored in reducing brain metastasis [61].

### Conclusions

This study is the first to report that Sdc1 expression on BC cells promotes brain metastasis. We propose a model in which Sdc1 fosters brain metastasis by facilitating BC cell transmigration specifically across the BBB and provide evidence for Sdc1-regulated cytokine/chemokine production and/or release from BC cells (Fig. 6) [62, 63]. We predict that Sdc1 promotes cancer cell BBB transmigration through a mechanism involving multiple cytokines/chemokines, and plan to further investigate this in future studies.

**Figure 6.**
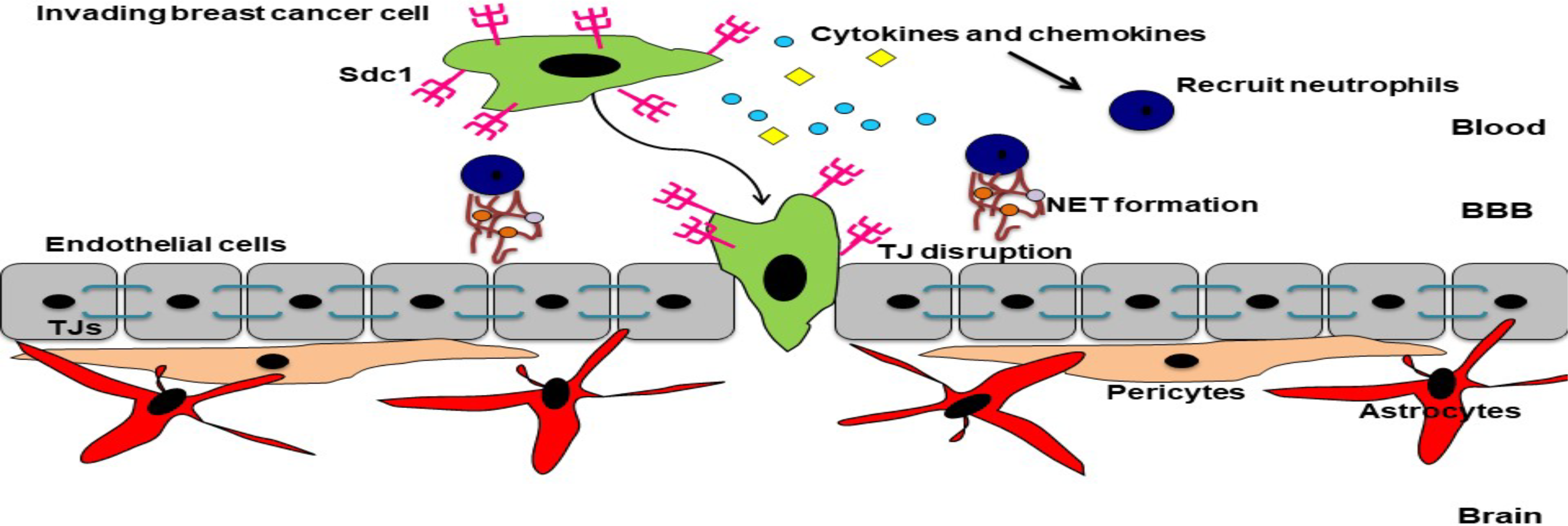
Working model. BC cells expressing Sdc1 produce and secrete high levels of various cytokines and chemokines. These factors are noted to play a role in BBB disruption. The chemokines can attract neutrophils and other leukocytes to facilitate transendothelial cell migration across the BBB. Neutrophils produce neutrophil extracellular traps (NETs), which induce BBB permeability. Overall, these factors may be key mediators of BC cell entry into the brain.

## Additional files

None.

## Declarations

None.

## Ethics approval and consent to participate

VCU IACUC Protocol AD10000943.

## Consent for publication

Not applicable.

## Competing interests

The authors declare that they have no competing interests.

## Availability of data and materials

All data and materials presented in this study are available from the corresponding author upon request.

## Funding

This work was financially supported by the American Cancer Society Research Scholar Grant ACS/RSG-123275-CSM.

## Authors’ contributions

Original idea and project development-JEK; acquisition of data-MRS, MP, NGV, SMW, MM, LM, CE, AA, SK, MS, MD, JS, MOI, JEK; analysis and interpretation of data-MRS, MP, NGV, SMW, MM, LM, CE, AA, SK, MS, MD, JS, MOI, JEK; writing-MRS, SMW, JEK. All authors have reviewed and approved this manuscript.

### Acknowledgements

We thank Megan Bliss-Morrow, David Finkelstein, Emily Lanning, Kaia Schwartz, Debra Chen, and Majid Jahromi for technical assistance with experiments, Azeddine Atfi for critical review of the manuscript, and Deborah Hurtado and Nikhail Mittal (ACEA Biosciences) for their guidance with the xCELLigence System. Flow cytometry and imaging work was performed in part at the Northwestern University Center for Advanced Microscopy and Flow Cytometry Core generously supported by NCI CCSG P30 CA060553 awarded to the Robert H. Lurie Comprehensive Cancer Center. Services in support of the research project were provided by the Virginia Commonwealth University Massey Cancer Center Flow Cytometry Core and Cancer Mouse Models Core Laboratory, supported, in part, with funding to the Massey Cancer Center from NIH-NCI Cancer Center Support Grant P30 CA016059. This work was supported by the American Cancer Society RSG-123275-CSM.

## Supplemental materials and methods

### Alteration of Sdc1 expression

As described previously, MDA-231 cells were infected with a lentivirus vector GINZEO (Open Biosystems) containing GFP and a shRNAmir to Sdc1 or a non-silencing (NS1) control sequence. Sdc1 KD was rescued by expression of mouse Sdc1 [31]. Sdc1 was amplified from pCMV6-Kan/Neo expression vector (Dharmacon) with primers containing EcoRI and SalI restriction sites and ligated into a pBABE retrovirus vector containing mCherry (gift from Dr. B. Parker, NU). This retrovirus vector was also used to overexpress Sdc1 in MDA-231 and 4T1 cells. An empty vector (EV) was infected into these cells as a control. All cells were selected twice by flow cytometry for the top 5% of GFP expression or mCherry expression. This is a heterogeneous population of cells that stably express the shRNAmir or Sdc1. Stable expression was determined by flow cytometry.

### Assessment of Sdc1 expression

For flow cytometry, cells were harvested with Versene (Invitrogen, 0.48mM EDTA) for 10 min at 37°C with gentle agitation. Cells were washed in PBS and resuspended in cold buffer containing 1% FCS. A total of 2 × 10^5^ cells per sample were used. Following centrifugation, cells were resuspended in Hanks’ balanced salt solution/2% BSA and incubated for 30 min at 4°C with titrated antibodies (20µg/ml mouse anti-Sdc 1 [B-A38, Bio-Rad]; rabbit anti-Sdc 2 [R&D], rabbit anti-Sdc 3 and 4 [ThermoFisher]), cells were then washed twice with PBS, incubated with secondary antibody for 30 min at 4ºC (1:100 dilution of donkey anti-mouse or rabbit-Cy5 [Jackson ImmunoResearch]), washed again twice with PBS and resuspended in 100 µl of PBS. The samples were analyzed using a Beckman Coulter FC500 flow cytometer. Human Sdc expression was analyzed comparing the relative amount of human Sdc stained cells with the IgG-controls and then KD cells were compared to NS1 cells. Isotype-matched antibodies were used as negative controls.

### Proliferation assay

Proliferation assays were performed as described [64] using MDA-231 infected with non-silencing (NS1) control or Sdc1 shRNAmir (Sdc1 KD) [31]. Briefly, 96-well plates were either coated with 50µl Cultrex BME (10mg/mL, Bio-Techne) for 30 min at 37°C. Cells (5,000/100µl/well) were seeded into each well with serum or serum-free media, and 2, 24, 48, 72, and 96-hours later cell growth was assessed after the addition of CellTiter-96-AQ_ueous_ One Solution Reagent according to manufacturer’s instructions (Promega). A one-way ANOVA was done to determine the statistical difference among sample means. The conservative Bonferroni’s multiple comparison post-test was combined with the ANOVA to compare differences between the mean values.

### Attachment of BC cells to endothelial and HA cells

iHUVEC and hCMEC cells (5 × 10^4^ cell/well) were attached to gelatin (0.2%, Sigma) coated coverslips and HA (5 × 10^4^ cells/well) were attached to poly-L-lysine (0.02mg/ml, Bio-Techne) coated coverslips in 24-well plates. Once the cells reached confluency (2-3 days), they were washed, incubated for 1 hour in serum-free media, and then either 2.5 × 10^5^ MDA-231 NS1 or Sdc1 KD cells were added to the wells in serum-free media. Each was performed in triplicate. After 30 min, unattached cells were washed off, cells in the wells were fixed with 4% paraformaldehyde, coverslips were mounted on slides with Prolong Diamond mounting medium (ThermoFisher) and 10 images per coverslip were taken using fluorescence microscopy. The number of tumor cells attached per image was counted using ImageJ software. Statistical analysis, unpaired Student’s t-test.

### Immunofluorescence

Briefly, frozen coronal brain sections were rehydrated with PBS, fixed (4% PFA, 30 minutes), blocked with 5% normal goat serum (90 minutes, Jackson ImmunoResearch), and incubated in PECAM-1 antibody (1:100; clone 2H8, gift from Dr. W. Muller, Northwestern University) overnight at 4ºC. After incubation with Cy3-conjugated goat anti-hamster secondary antibody (1:100, 1 hour, Jackson ImmunoResearch) and DAPI (167ug/ml), slides were cover-slipped with Prolong Diamond mounting medium (ThermoFisher).

### Cytokine array and validation

The human cytokine array kit (R&D Systems) was used according to the manufacturer’s instructions. In brief, 24-hour CM from MDA-231 NS1 and Sdc1 KD cells was collected and applied to membranes overnight, signals were detected after incubation with antibody cocktails and streptavidin–HRP followed by chemiluminescent detection. GE ImageQuant LAS 4000 software was used for quantification of the spots. Validation was performed using Bio-plex Multiplex Immunoassay System (Bio-Rad) according to manufacturer’s instructions. Statistical analysis, one-way ANOVA with Bonferroni’s multiple comparison post-test.

### Statistical analysis

Each sample was performed in triplicate or quadruplicate, and the experiments were repeated at least three times. Data were imported into GraphPad Prism Software for statistical analysis and reported as mean ± SEM. Analysis was performed using the appropriate statistical methods as indicated in the methods. *p* < 0.05 was considered significant.

## Supplemental Figure legends

**Supplemental Figure 1.**
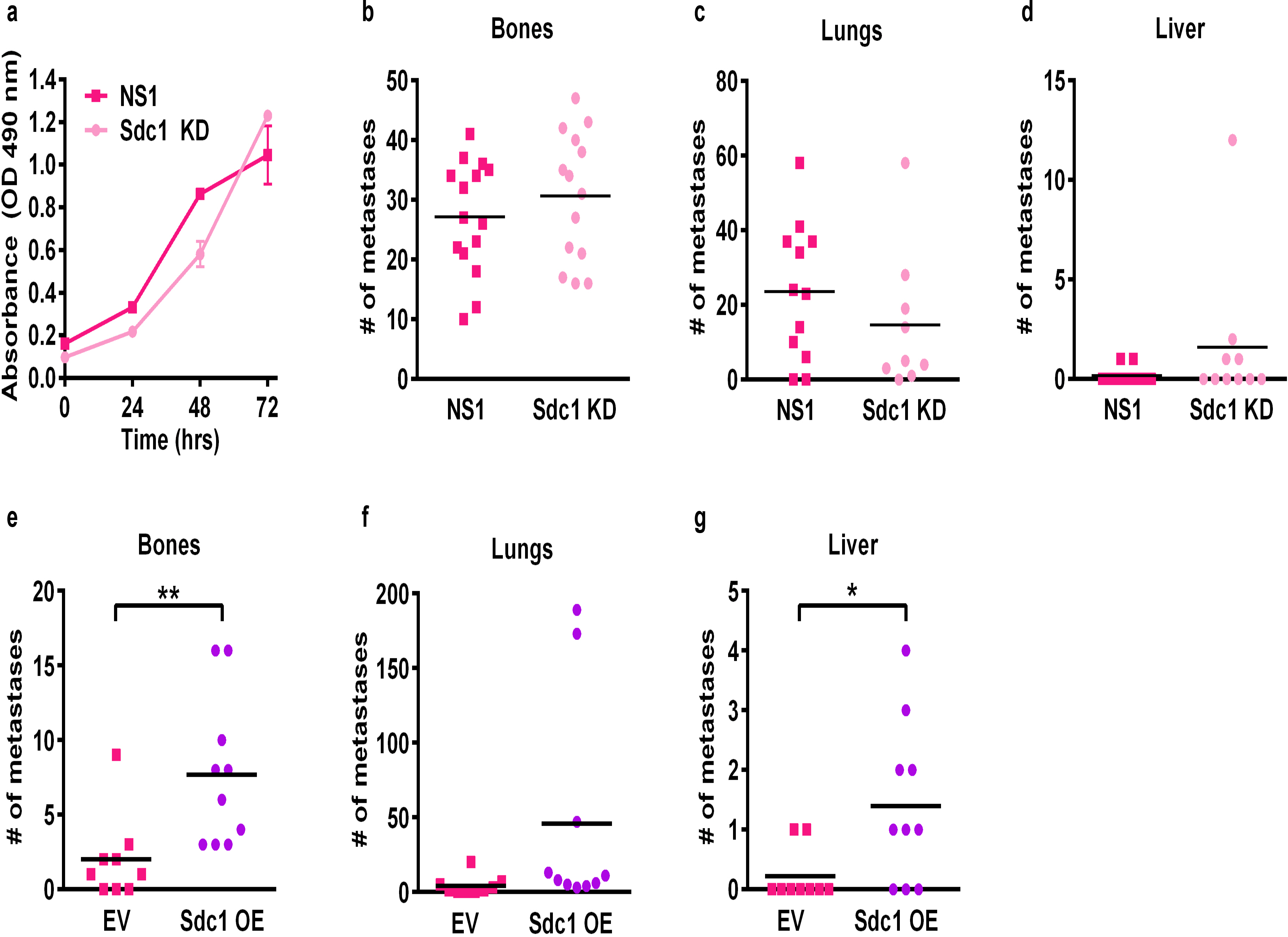
The effects of silencing Sdc1 in BC cells on other organs. **a** Silencing of Sdc1 expression does not affect proliferation of MDA-231 cells grown in serum-containing media on BME/Matrigel. **b-d** Decreased expression of Sdc1 in MDA-231 had no effect on (b) bone, (c) lung, or (d) liver metastases after intracardiac injection of these cells compared to control NS1 cells. **e-g** Overexpression of Sdc1 in 4T1 cells had no effect on (f) lung metastases but significantly increased metastases to the bones (e) and liver (g) compared to the control EV cells. *p<0.05, **p<0.01.

**Supplemental Figure 2.**
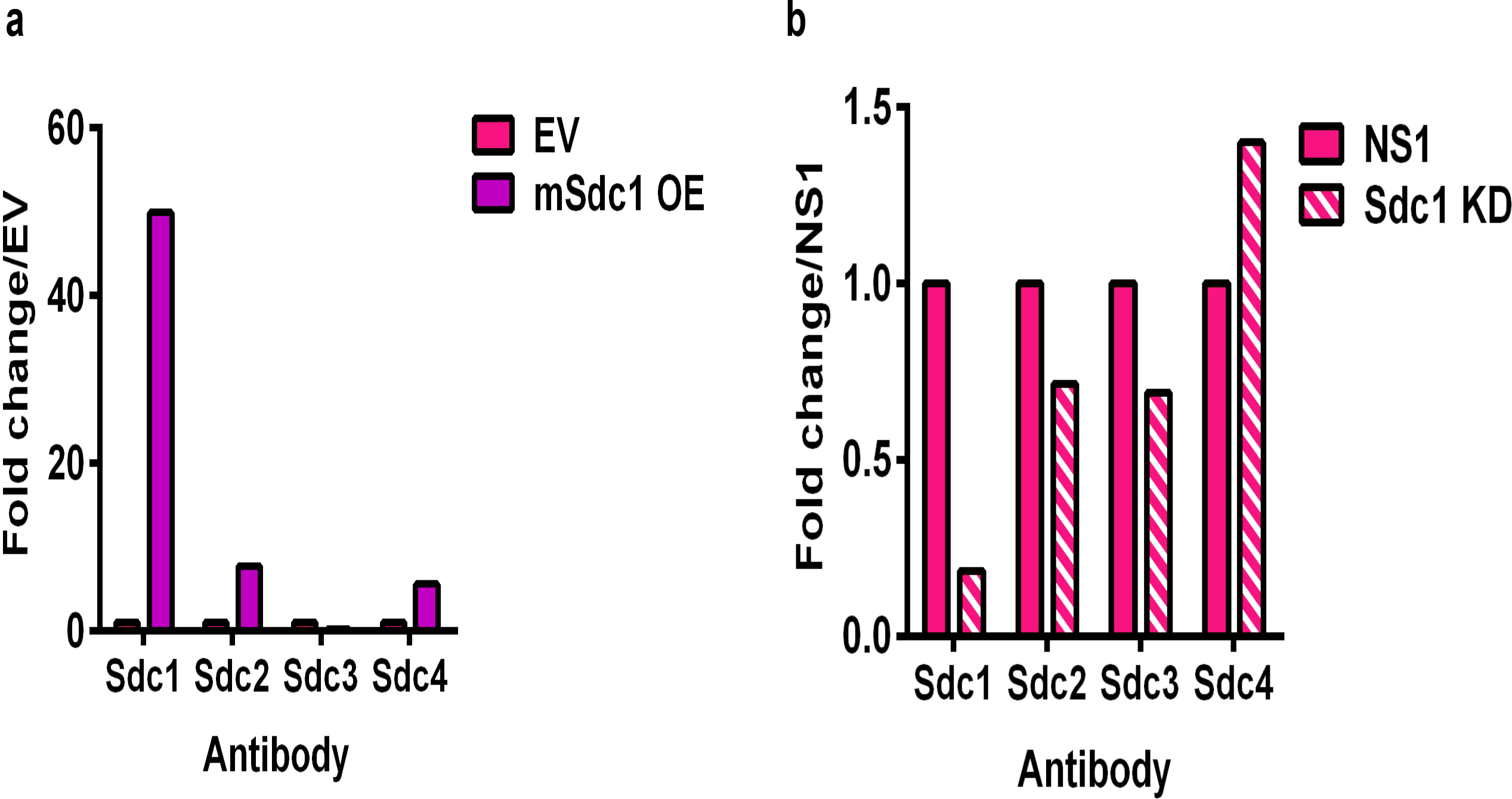
Expression of Sdc1 is increased by 50-fold in 4T1 cells and decreased by 5-fold in BT549 cells. Flow cytometry was used to measure the surface expression of the Sdcs in **a** 4T1 after overexpression of Sdc1 and **b** BT549 cells after silencing of Sdc1. Bars, mean.

**Supplemental Figure 3.**
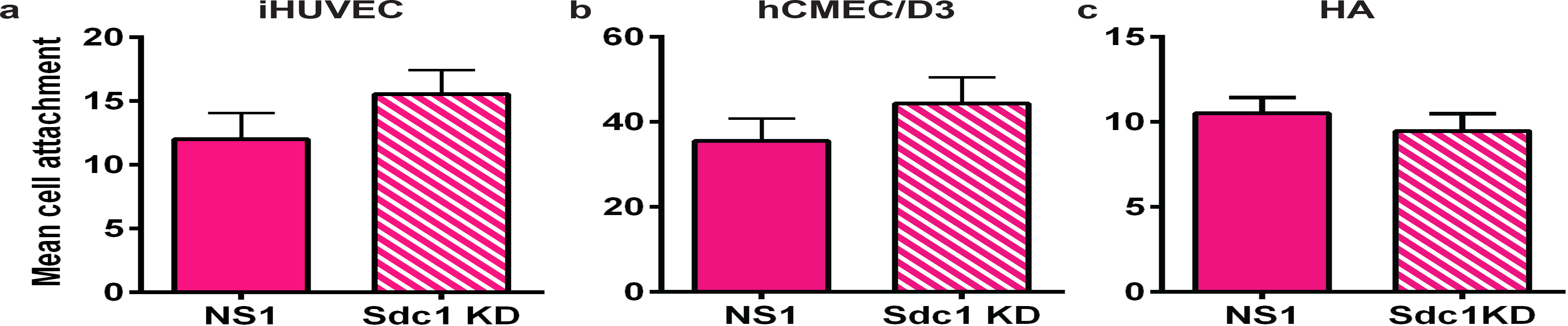
Silencing of Sdc1 expression has no effect on in vitro cell attachment to the endothelium or astrocytes. There is no difference in MDA-231 NS1 and Sdc1 KD cell attachment to iHUVEC (**a**), hCMEC (**b**), or HA (**c**). n=3, Bars, mean cell attachment averaged/image, 10 images/well, 3 wells/group ± SEM.

**Supplemental Figure 4.**
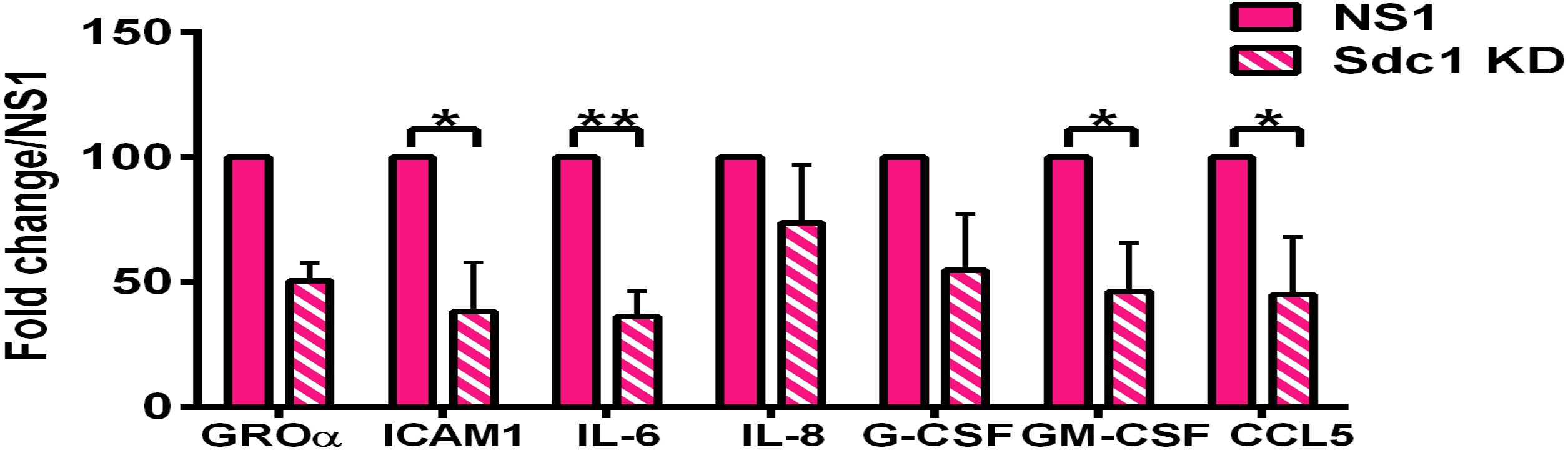
Silencing Sdc1 affects cytokine expression in MDA-231 cells. Quantification of cytokine array (R&D Systems) incubated with 24-hour CM from MDA-231 NS1 and Sdc1 KD cells. n=3, *p<0.05, **p<0.01.

## Abbreviations

BBB: blood-brain barrier
Sdc1: syndecan-1
MDA-231: MDA-MB-231
Sdc1 KD: syndecan-1 knock-down
NS1: non-silencing sequence
Sdc1 OE: syndecan-1 over-expression
EV: empty vector
HER2+: Human epidermal growth factor receptor 2 positive
TNBC: triple-negative breast cancer
DMEM: Dulbecco’s modified Eagle’s medium
FBS: fetal bovine serum
BME: basement membrane extract
NSG: NOD.Cg-*Prkdc^scid^ Il2rg^tm1Wjl^*/SzJ
ANOVA: analysis of variance
iHUVEC: immortalized human umbilical vein endothelial cells
HA: human astrocytes
CI: cell index
PBS: phosphate-buffered saline
PFA: paraformaldehyde
DAPI: 4’,6-Diamidino-2-phenylindole
GFP: green fluorescent protein
PECAM-1: platelet endothelial cell adhesion molecule-1
CM: conditioned medium
GRO-ɑ: growth regulated oncogene-alpha
ICAM1: intercellular adhesion molecule 1
IL-8: Interleukin-8
IL-6: Interleukin-6
G-CSF: granulocyte colony-stimulating factor
GM-CSF: granulocyte-macrophage colony-stimulating factor
CCL5: C-C motif chemokine ligand 5
TMA: tissue microarray
TCGA: The Cancer Genome Atlas.

